# A Biphasic Effect of Alcohol on Endothelial Plasticity Through Regulation of Endothelial-to-Mesenchymal Transition

**DOI:** 10.64898/2026.04.14.718463

**Authors:** Weimin Liu, Yusof Gusti, Fathima Athar, Naresh K Rajendran, Paul A. Cahill, Eileen M. Redmond

## Abstract

**Background:** Alcohol consumption influences cardiovascular disease, but whether it does so by affecting endothelial plasticity is unknown. We tested whether alcohol regulates endothelial-to-mesenchymal transition (EndMT) to influence arterial pathology.

**Methods:** HCAEC and HUVEC were exposed to inflammatory cytokines (TGFβ ± IL1β) or hypoxia in the presence of ethanol (0-100 mM). EndMT was assessed by changes in cell marker expression, SNAIL levels, and migration assays. In vivo, carotid ligation was performed in mice gavaged with/without either daily moderate ethanol (2-drink equivalent/d) or episodic binge exposure (7-drink equivalent, 2 days/week) and myo-endothelial cell population assessed.

**Results:** Cytokines and hypoxia induced EndMT in vitro, characterized by loss of endothelial markers, increased mesenchymal markers, elevated SNAIL, and enhanced migratory capacity. Low-to-moderate dose ethanol (5–25 mM) attenuated these changes, preserving endothelial phenotype, whereas high dose ethanol (50–100 mM) either had no effect or exacerbated EndMT. The inhibitory effect of moderate ethanol on cytokine- and hypoxia-induced changes in αSMA and Cdh5 expression was abrogated by γ-secretase inhibition, consistent with involvement of Notch signaling. Carotid ligation induced neointimal formation and accumulation of myo-endothelial cells indicative of EndMT. Daily moderate ethanol significantly attenuated neointimal hyperplasia and diminished the myo-endothelial cell population, whereas in contrast, episodic binge ethanol exposure increased pathologic remodeling and myo-endothelial cell abundance.

**Conclusions:** Alcohol modulates endothelial trans-differentiation in a biphasic manner. Low-to-moderate alcohol exposure suppresses EndMT and limits pathological remodeling, whereas binge-level exposure promotes these processes. These findings identify regulation of endothelial plasticity as a potential novel mechanism linking alcohol consumption patterns to vascular disease risk.

**NEW AND NOTEWORTHY:** We identify a previously unrecognized biphasic effect of alcohol on endothelial phenotypic plasticity. Low-to-moderate dose alcohol suppresses endothelial-to-mesenchymal transition (EndMT), whereas high-level (binge) exposure promotes this pro-atherogenic process. Given the central role of EndMT in vascular remodelling and atherosclerosis, these findings provide a mechanistic framework linking alcohol consumption patterns and cardiovascular disease risk – potentially explaining both the protective effect at low/moderate levels, and the detrimental impact of heavy alcohol use.

**Graphical abstract:** 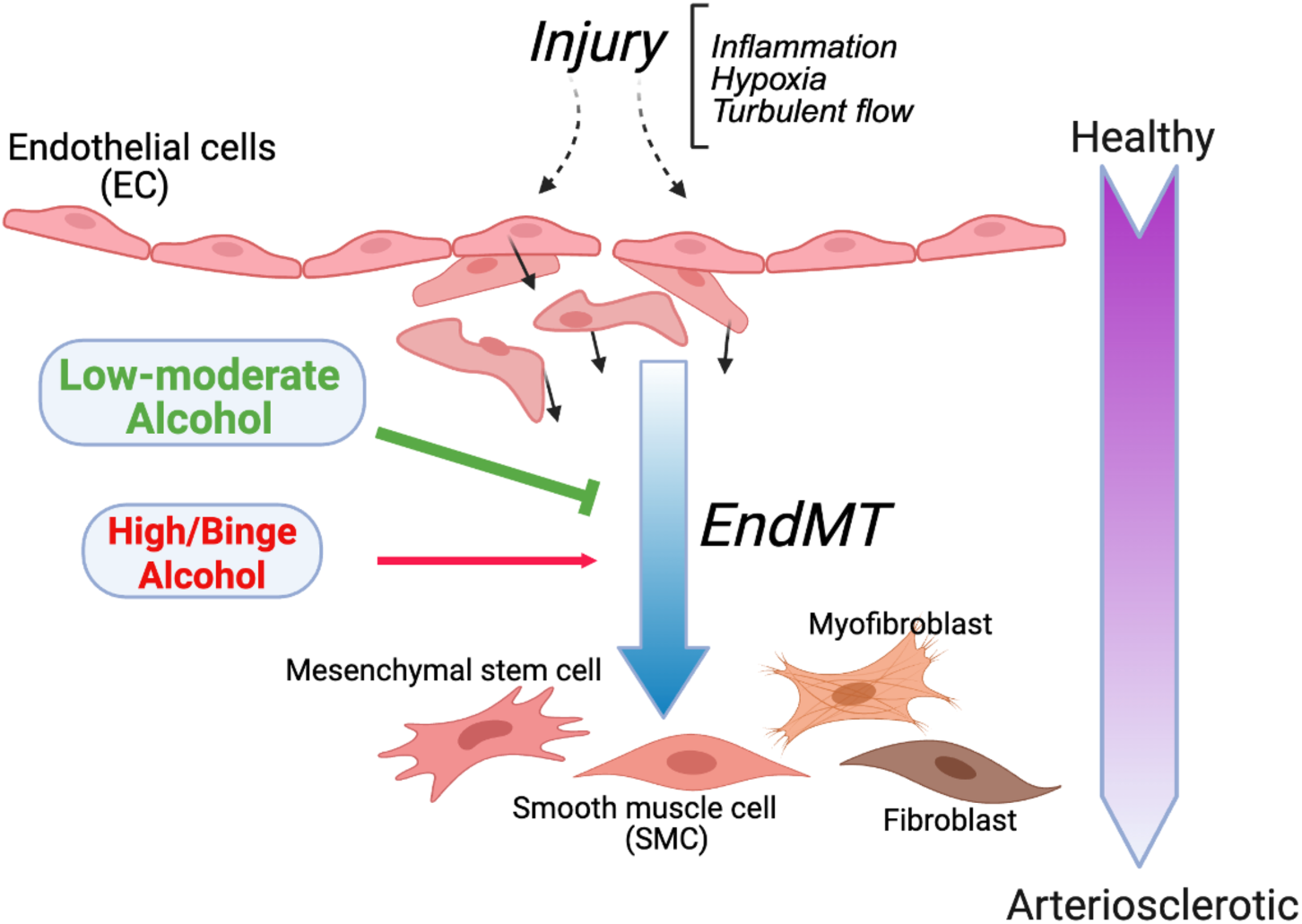

Injurious stimuli can trigger endothelial cells (EC) to undergo endothelial-to-mesenchymal transition (EndMT) that contributes to arterial remodeling and disease. EndMT is regulated in a biphasic manner by alcohol with low-to-moderate levels (1-3 drink equivalent) suppressing EndMT and attenuating vascular remodeling, whereas higher level/binge exposure (7 drink equivalent) promotes these processes. *Graphic created using Biorender*.

## INTRODUCTION

Alcohol (ethanol) consumption is associated with a variety of health effects(10, 15). In the context of atherosclerotic cardiovascular disease (ASCVD) that causes myocardial ischemia and stroke, epidemiologic studies support a biphasic or ‘J-shaped’ relationship, whereby low-to-moderate alcohol consumption (generally 1–3 drinks/day; blood alcohol concentration [BAC] ∼5–25 mM) is associated with reduced risk, while higher levels of consumption (BAC ≥50 mM), particularly in binge patterns, are associated with increased morbidity and mortality(7, 24, 28). Understanding the cellular and molecular mechanisms mediating these complex dose-dependent effects of alcohol on ASCVD remains an important and unresolved question.

Endothelial-to-mesenchymal transition (EndMT) is a dynamic process wherein endothelial cells lose their endothelial characteristics and acquire mesenchymal features, including smooth muscle-like and myofibroblast-like phenotypes. EndMT is essential during embryonic cardiovascular development for valve formation and septation (35), but its dysregulation has been increasingly implicated in adult pathologies such as fibrosis, cancer, and especially cardiovascular disease including atherosclerosis and hypertension (9, 16). In atherogenesis, EndMT represents a continuum of phenotypic and functional changes in which endothelial cells lose cell–cell junctions and barrier integrity, degrade the basement membrane, and acquire migratory and proliferative capacity, enabling their accumulation within the sub-intimal space, thus, contributing directly to neointimal hyperplasia and vessel remodeling (11, 17). While in general the regulation of EndMT remains poorly understood, available evidence suggests that it is predominantly driven by cytokines such as TGF-β, TNF-α and IL-1β, as well as physical and environmental stressors such as disturbed shear stress and hypoxia, respectively. Signaling pathways implicated in EndMT include WNT, Smad, and Notch, with pathways converging to upregulate transcription factors such as Snail, Slug, Twist and Zeb1/2 (17) (1).

Endothelial cell function is a critical determinant of vascular homeostasis (4) and alcohol’s impact on these cells has been previously interrogated. A biphasic effect of alcohol on several key endothelial functions including barrier function, vasoactive agent production, inflammatory cytokine release, and cell adhesion molecule expression, and on atherosclerotic pathology has been previously demonstrated (14, 21, 25). These studies predict low-to-moderate dose alcohol to be anti-atherogenic, with higher level exposures pro-atherogenic (14). However, whether alcohol directly influences endothelial phenotypic plasticity - specifically EndMT - remains unknown.

Accordingly, the aim of this study was to determine whether alcohol modulates EndMT in a dose- and pattern-dependent manner to influence pathologic vascular remodeling. Using complementary in vitro models of cytokine- and hypoxia-induced EndMT in human endothelial cells and an in vivo endothelial lineage–traced carotid ligation model, we demonstrate that low-to-moderate alcohol exposure suppresses EndMT-like phenotypic switching and attenuates neointimal formation, whereas binge-level exposure promotes these processes. These findings identify regulation of endothelial plasticity as a potential mechanistic link between alcohol consumption patterns and vascular disease outcomes.

## MATERIALS AND METHODS

### Cell culture and treatments

Human coronary artery endothelial cells (HCAEC, C-12221) and Human umbilical vein endothelial cells (HUVEC-pooled, C-12203) were obtained from PromoCell. Cells were cultured on fibronectin-coated (5 µg/mL in PBS, Corning^TM^) plates in either endothelial cell growth medium (C-22010, Promocell) or EBM-2 basal media with supplements and growth factors (EMB-2 growth media) (Lonza Walkersville CC3156 and CC4176). Cells from passages 5-12 were used for all experiments. Cells were pretreated with ethanol (5-100 mM) for 90 min prior to other treatments. EndMT was induced by treating cells with TGFβ1 (10 ng/mL, Cat. # 100-21, PeproTech), a combination of IL1β (5 ng/mL, Cat. # 200-01B, PeproTech) and TGFβ2 (10 ng/mL, 100-35B, PeproTech), or hypoxia (5% O_2_). TGFβ1 treatment was for either 2, 4, or 7 days, while IL1β+TGFβ2 treatment was for 2 days. For hypoxia experiments, cells in EBM2 growth media with supplements and 1 % FBS were placed in a hypoxic incubator chamber (Stemcell Technologies) purged with hypoxic mixed gas every day, for 4 days. Where indicated, cells were treated with Notch inhibitor DAPT (20 μM, Cat. #2634, Tocris,) or Notch ligand DLL4 (2 μg/ml, Cat. # 1506-D4, R&D Systems). Media with reagents was replenished every alternate day.

### Real time qPCR

Total RNA was isolated using ReliaPrep TM RNA cell Miniprep System (Cat. # Z6011, Promega) following manufacturer’s instructions. DNase-treated RNA was either used in one-step qRT-PCR (Cat. # 208354, QuantiNova RT-PCR kit) or for cDNA synthesis (1725038, Biorad). qRT-PCR was performed using gene specific primers and QuantStudio 3 (Applied Biosytems). List of primers is provided (Table 1).

**Table 1.**
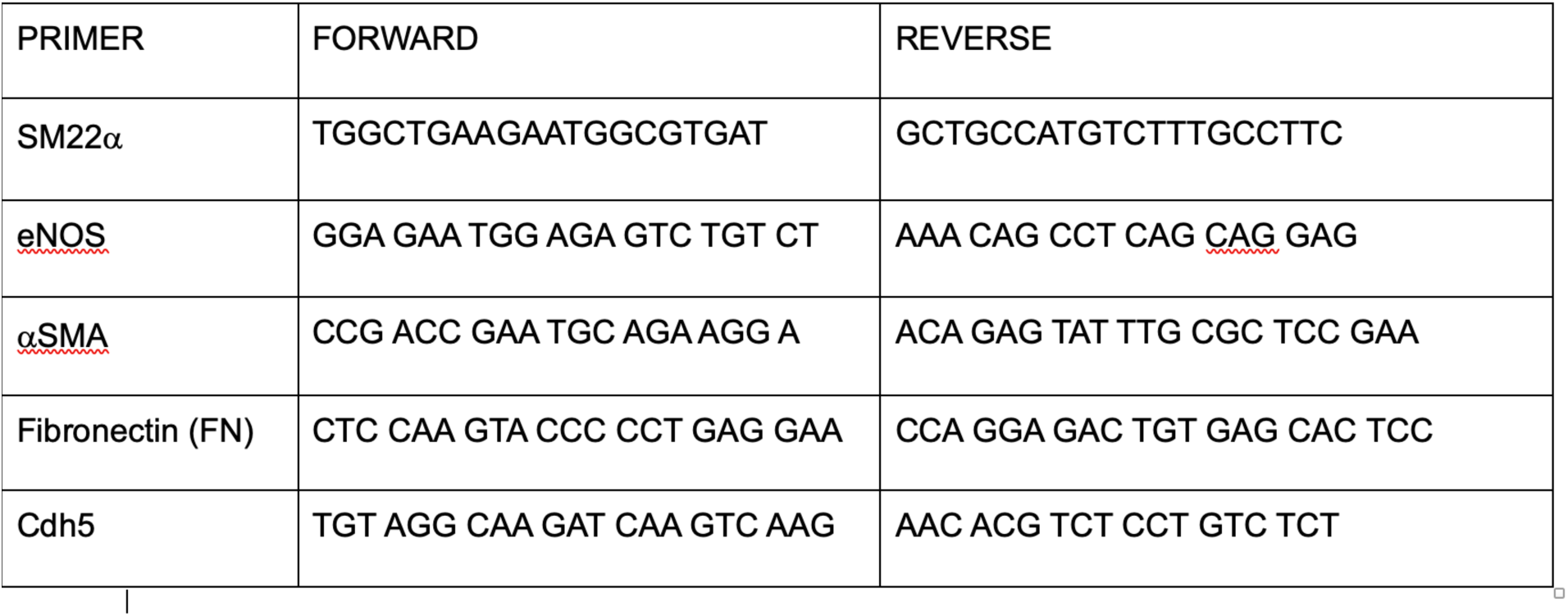

#### Western blotting

Cells were washed twice with PBS and lysed in RIPA buffer (Cat. # 89901, Thermo Scientific) and protein was estimated using Pierce™ BCA Protein Assay Kit (Cat. # A55865, Thermo Scientific). Lysates were electrophoresed and transferred onto PVDF membranes. Blots were blocked using 5% Blotting-Grade Blocker (Cat. # 1706404, Biorad) or BSA (Cat. # A7030, Sigma), depending on the antibody, in Tris-buffered saline with 0.1 % Tween-20 (TBST). Blots were then incubated in 1/10^th^ blocking buffer overnight at 4° C with primary antibody at a dilution recommended by the manufacturer. Following 3X washes with TBST, 5 min each, blots were incubated with HRP-conjugated secondary antibody (Cat. # 7074, 7076, CST) in 1/10^th^ blocking buffer at 1:2000 dilution for 1-2 h at room temperature (RT). Bands were visualized by chemiluminescence using ECL reagent (Cat. # 1705060, Biorad) and imaged using ChemiDoc XR Imaging System (Bio-Rad). Primary antibodies used were human CD31 (Cat. # 3528, Cell Signaling Technology (CST)), CDH5 (Cat. # 2500, CST), SM22α (Cat. # 36090, CST), SNAIL (Cat. # 3879, CST), GAPDH (Cat. # MA5-15738, Invitrogen), and β-Actin (Cat. # 4979, CST).

#### Immunostaining

Cells grown on fibronectin-coated coverslips were fixed in 10% formaldehyde in PBS (10 min at RT), washed 3X with PBS, permeabilized with 0.5 % Triton-X100 in PBS (6 min, RT), washed 3X, and blocked with 0.5 % fish gelatin in PBS (1-2 h). Cells were incubated with primary antibody in 1/10^th^ gelatin in PBS for 2-3 h. Following 3X washes, cells were incubated with flourescently-labelled secondary antibody for 1-2 h. Following 3x washes cells were mounted on slides using Vectashield Plus Antifade mount medium with DAPI (Cat. # H20002, Vector Laboratories) and scanned with confocal microscope (Nikon A1R HD, Pikachu). Images were analyzed using Image J. Primary antibodies used include human CD31 (Cat. # AF806, R&D Systems), αSMA (MAB1420, R&D systems), SM22α (Cat. # 36090 CST), and CDH5 (Cat. # 2500, CST).

Secondary antibodies used were Sheep IgG NorthernLights™ NL557 (Cat. # NL010, R&D Systems), Mouse IgG NorthernLights™ NL557 (Cat. # NL007, R&D Systems), and Anti-rabbit IgG Alexa Fluor® 488 Conjugate (Cat. # 4412, CST).

### Migration assay

Following treatment, 30,000-50,000 cells were seeded in 3-well culture inserts (Cat. # 80369, Ibidi) for at least 12-24 h in their respective treatment media. The inserts were then removed, 1-2 ml of respective media added, and cell migration was imaged at regular intervals for 12 h. Images were analyzed using Image J and a plugin for scratch assay(30).

### Mice

Mice expressing a tamoxifen-inducible Cre recombinase under the control of the vascular endothelial cadherin (Cdh5(PAC)) promoter i.e., C57Bl/6-Tg(Cdh5-cre/ERT2)1Rha (Taconic Biosciences, #13073) were crossed with Ai9 reporter mice (B6.Cg-Gt(ROSA)^26Sortm9(CAG-tdTomato)Hze^/J (Jax Labs, #007909) to generate both male and female Cdh5-Ai9 transgenics used in carotid ligation experiments.

### Tamoxifen treatment

To induce Cre-LoxP recombination and permanently label endothelial cells, Cdh5-Ai9 mice (average weight 20g, 6-8 weeks old) were injected with tamoxifen, 75 mg/kg IP, for 5 consecutive days. They were then rested for 1 week (during which time they received, or not, EtOH treatments) before ligation surgery or sham operation. tdTomato expression in these mice reflects lineage labeling of Cdh5-expressing endothelial cells.

### Ethanol (EtOH) treatments

Daily moderate ethanol: Mice received ethanol by oral gavage at 0.8 g/kg/day (200 proof ethanol, ACS/USP grade; maximum volume 200 μL), corresponding to approximately 2 drinks/day and a peak blood alcohol concentration (BAC) of ∼15 mM (0.07%), as previously described (13). Treatment was initiated 1 week prior to ligation, paused on the day of surgery, and resumed 1 day post-surgery, continuing daily for up to 2 weeks until tissue harvest. Control animals received a calorically matched water–cornstarch solution. Episodic binge ethanol: Mice received ethanol at 2.8 g/kg/day on 2 consecutive days per week, corresponding to ∼7 drinks/day and a peak BAC of ∼50 mM (0.2%) (21). This regimen was administered 1 week prior to ligation and continued for 2 weeks post-surgery. Both ethanol groups received equivalent total weekly ethanol exposure (5.6 g/kg/week).

### Mouse carotid ligation or sham surgery

Left common carotid artery ligation was performed as previously described (13, 20). Briefly, 1 week after tamoxifen-induced recombination, mice were anesthetized with inhaled isoflurane and placed on a temperature-controlled surgical platform. Preoperative analgesia was administered using sustained-release buprenorphine (0.5–1.0 mg/kg, subcutaneous), with repeat dosing every 72 hours as needed. Following a midline cervical incision, the left common carotid artery was exposed under a dissecting microscope and ligated just proximal to the bifurcation using 6-0 silk suture. In sham-operated mice, the artery was exposed but not ligated. The incision was closed in two layers (muscle and skin), and animals were allowed to recover under observation. All procedures were approved by the University of Rochester Animal Care Committee and conformed to NIH guidelines for the care and use of laboratory animals.

### Histomorphometric analysis

Two weeks post-ligation, mice were anesthetized (ketamine/xylazine) and perfusion-fixed with 4% paraformaldehyde in sodium phosphate buffer (pH 7.0). Carotid arteries were harvested, paraffin-embedded, and sectioned. Serial cross-sections (5 μm thickness; 10 sections per level) were obtained every 200 μm over a 2 mm segment distal to the carotid bifurcation. Sections were stained with Verhoeff–Van Gieson stain to visualize elastic laminae and imaged using a Nikon TE300 microscope equipped with a Spot RT digital camera. Morphometric analysis was performed using ImageJ2 (v2.14.0/1.54i). Lumen area was calculated from luminal circumference assuming circular geometry. Intimal area was defined as the region between the lumen and internal elastic lamina (IEL), medial area between the IEL and external elastic lamina (EEL), and adventitial area between the EEL and outer vessel boundary, as previously described (20, 23).

#### Immunohistochemistry and analysis of fluorescently labeled cells

Carotid sections were stained with anti–α-smooth muscle actin (αSMA; Invitrogen, MA5-11547, 1:200) followed by Alexa Fluor 647–conjugated goat anti-mouse secondary antibody (Abcam, ab150115), and anti-tdTomato (Abcam, ab62341, 1:200) followed by Alexa Fluor 594–conjugated goat anti-rabbit secondary antibody (Invitrogen, A11013). Antigen retrieval was performed in 10 mM sodium citrate buffer (pH 6.0) at near-boiling temperature for 10 minutes, followed by cooling for 30 minutes. Sections were washed in deionized water and PBS prior to imaging. Images were acquired using an Olympus FV1000 or Nikon A1R HD confocal microscope. Quantification of fluorescently labeled cells was performed using QuPath (v0.5.1) using automated cell detection and object classification functions. Cells were categorized as endothelial (tdTomato^+^/Cdh5^+^), mesenchymal (αSMA^+^), or double-positive myoendothelial (Cdh5^+^αSMA^+^), and quantified across entire cross-sections and neointimal compartments.

#### Statistical analysis

Data are presented as mean ± SEM. In vitro experiments were performed in at least duplicate with a minimum of three independent biological replicates. For in vivo studies, n = 10 mice per group (5 males, 5 females) unless otherwise stated. Where indicated, analyses were performed separately by sex and combined. Because vascular remodeling was more pronounced in males, immunohistochemical analyses of double-positive (myoendothelial) cells were performed in male mice to maximize sensitivity. Statistical comparisons between two groups were performed using unpaired Student’s t-test with Welch correction. Multiple group comparisons were analyzed by one-way ANOVA followed by Tukey’s or Dunnett’s post hoc tests (GraphPad Prism v10.6.1). A p-value < 0.05 was considered statistically significant.

## RESULTS

### Biphasic effect of ethanol on cytokine- and hypoxia-induced EndMT in cultured human endothelial cells

Human coronary artery endothelial cells (HCAEC) were exposed to inflammatory cytokines (TGFβ1, IL-1β) or hypoxia in the presence or absence of ethanol (5–100 mM). Endothelial-to-mesenchymal transition (EndMT) was assessed by expression of endothelial markers (eNOS, CD31/PECAM-1, Cdh5), mesenchymal markers (αSMA, SM22α), the extracellular matrix protein fibronectin (FN), and the transcription factor SNAIL. Cytokine and hypoxic stimulation induced a robust EndMT-like response, characterized by co-ordinated downregulation of endothelial markers and upregulation of mesenchymal markers, FN, and SNAIL at mRNA and/or protein levels (Fig 1). Co-treatment with low-to-moderate ethanol concentrations (5–25 mM) significantly attenuated these changes, preserving endothelial marker expression while suppressing mesenchymal and transcriptional EndMT-associated responses. In contrast, higher ethanol concentrations (50–100 mM) failed to inhibit EndMT and, in some instances, showed a trend toward enhancement of the response. These findings demonstrate a dose-dependent, biphasic effect of ethanol on endothelial phenotypic switching.

**Fig 1.**
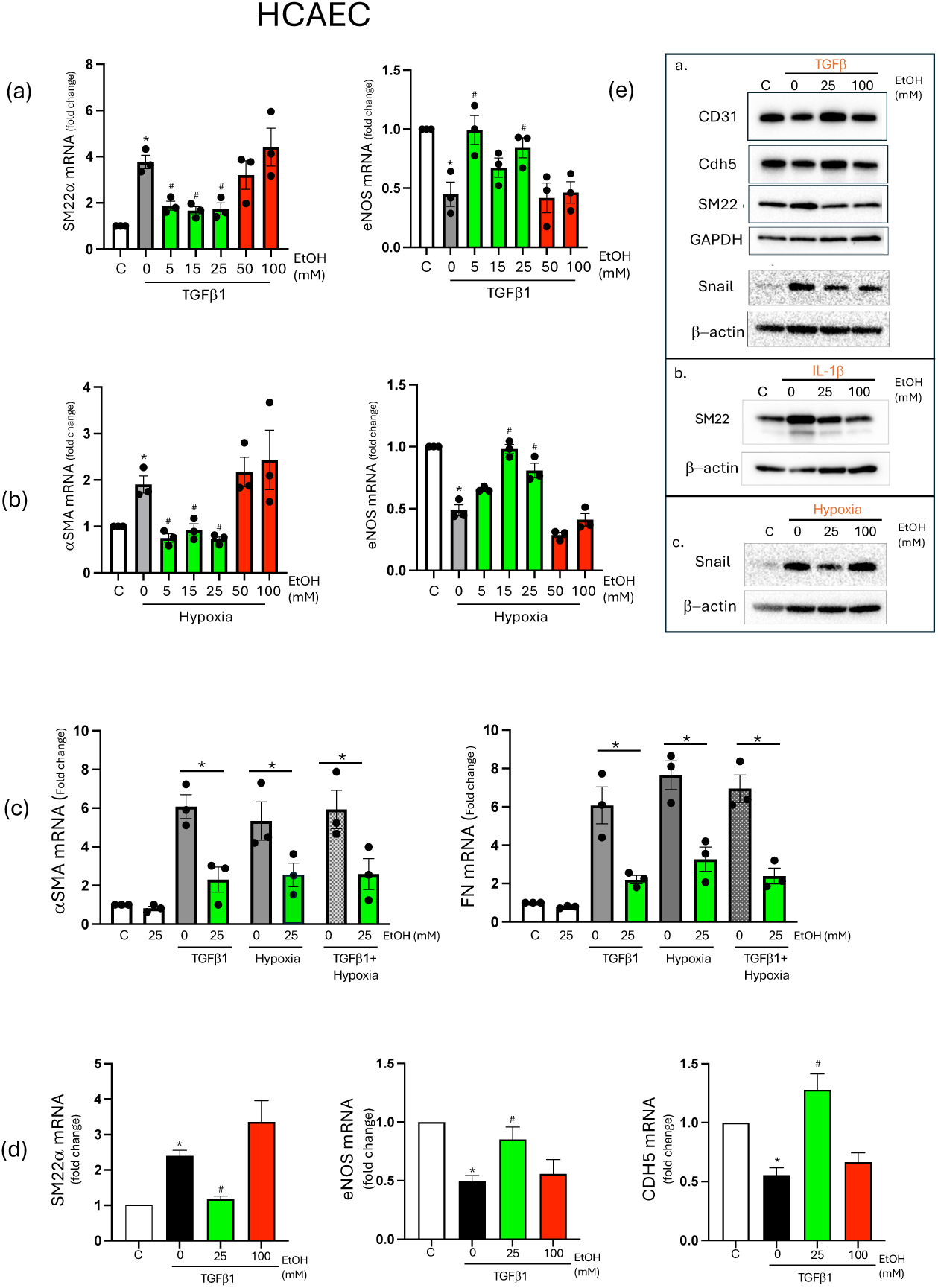
Biphasic effect of ethanol on cytokine and hypoxia-induced EndMT in cultured endothelial cells. (a-c) SM22α, eNOS, αSMA, and Fibronectin (FN) mRNA levels determined by qRT-PCR in human coronary artery endothelial cells (HCAEC) following treatment with TGFβ1 (10 ng/ml, 7d), hypoxia (5% O_2_, 4d) or TGFβ1 plus hypoxia (4d), in the absence or presence of ethanol (EtOH) at concentrations indicated (0-100 mM range). (d) SM22α, eNOS, and Cdh5 mRNA levels in HCAEC treated with TGFβ1 (10 mg/ml, 2d) +/- EtOH 25 mM or 100 mM. (e) Representative western blots showing CD31, Cdh5, SM22α, and SNAIL protein expression in HCAEC treated with TGFβ1, IL-1β, or Hypoxia +/- EtOH (25 mM or 100 mM). β-actin or GAPDH were used as loading controls. Data are mean±SEM, n=3. *p<0.05 vs control (no treatment), #p<0.05 vs TGFβ1 or hypoxia.

### Differential effect of moderate and high-dose ethanol on EndMT in HCAEC and HUVEC; immunocytochemical analysis

To confirm these findings at the cellular level, immunofluorescence analysis was performed in both HCAEC and human umbilical vein endothelial cells (HUVEC). Treatment with cytokines (TGFβ +/-IL-1β), or exposure to hypoxia, decreased CD31 expression and increased αSMA and SM22α expression consistent with EndMT-like phenotypic switching (Fig 2). These changes were markedly attenuated in cells treated with moderate dose EtOH (25 mM), whereas high-dose ethanol (100 mM) did not confer protection (Fig 2). The consistency of these effects across two endothelial cell types supports a generalizable modulatory effect of ethanol on endothelial phenotype.

**Fig 2.**
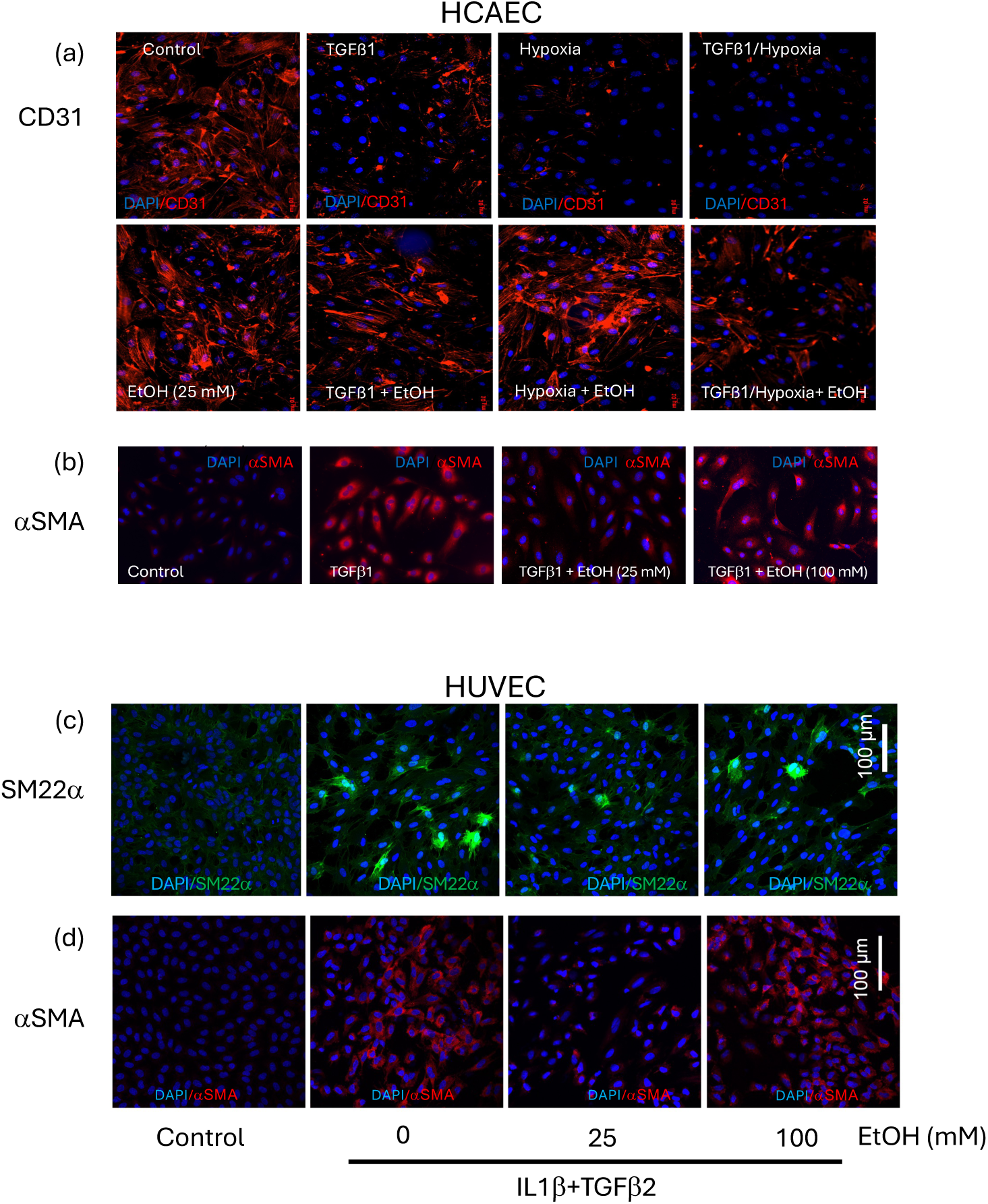
Differential effect of moderate and high dose ethanol on EndMT in HCAEC and HUVEC determined by immunocytochemistry. (a) CD31 expression in HCAEC treated with TGFβ1 (10ng/ml), Hypoxia (5% O_2_), or TGFβ1+ Hypoxia in the absence or presence of moderate dose ethanol (EtOH 25 mM). (b) αSMA expression in HCAEC treated with TGFβ +/- either moderate dose ethanol (25 mM EtOH) or high dose ethanol (100 mM EtOH). (c) SM22α and (d) αSMA expression in HUVEC treated with IL1β+TGFβ2 in the absence or presence of either 25 mM EtOH or 100 mM EtOH as indicated. Representative immunofluorescence images, from at least 3 experiments, shown.

### Attenuation of cytokine- and hypoxia-induced changes in αSMA and Cdh5 expression in HCAEC by moderate ethanol is Notch signaling-dependent

To explore potential mechanisms, αSMA expression was assessed in HCAEC exposed to TGFβ or hypoxia in the absence or presence of ethanol (25 mM or 100 mM), with or without the γ-secretase inhibitor DAPT. Moderate dose ethanol (25 mM) significantly attenuated TGFβ- and hypoxia-induced αSMA expression, effects that were abolished by DAPT (Fig 3a), consistent with involvement of Notch signaling. In contrast, high dose ethanol (100 mM) further increased TGFβ- induced αSMA expression, an effect that was unaffected by DAPT. In a manner similar to moderate dose ethanol, activation of Notch signaling using the ligand DLL4 suppressed TGFβ- induced αSMA expression, an effect reduced by DAPT (Fig 3b). Moreover, moderate dose ethanol (25 mM), but not high dose (100 mM), prevented cytokine-induced decreases in Cdh5 expression in HCAEC determined by immunocytochemistry (Fig 4). This effect of moderate ethanol was blocked by DAPT (Fig 4). Together, these findings suggest that Notch signaling contributes to the inhibitory effects of moderate ethanol on EndMT-like responses, whereas the stimulatory effects of higher ethanol exposure when present appear to be Notch-independent.

**Fig 3.**
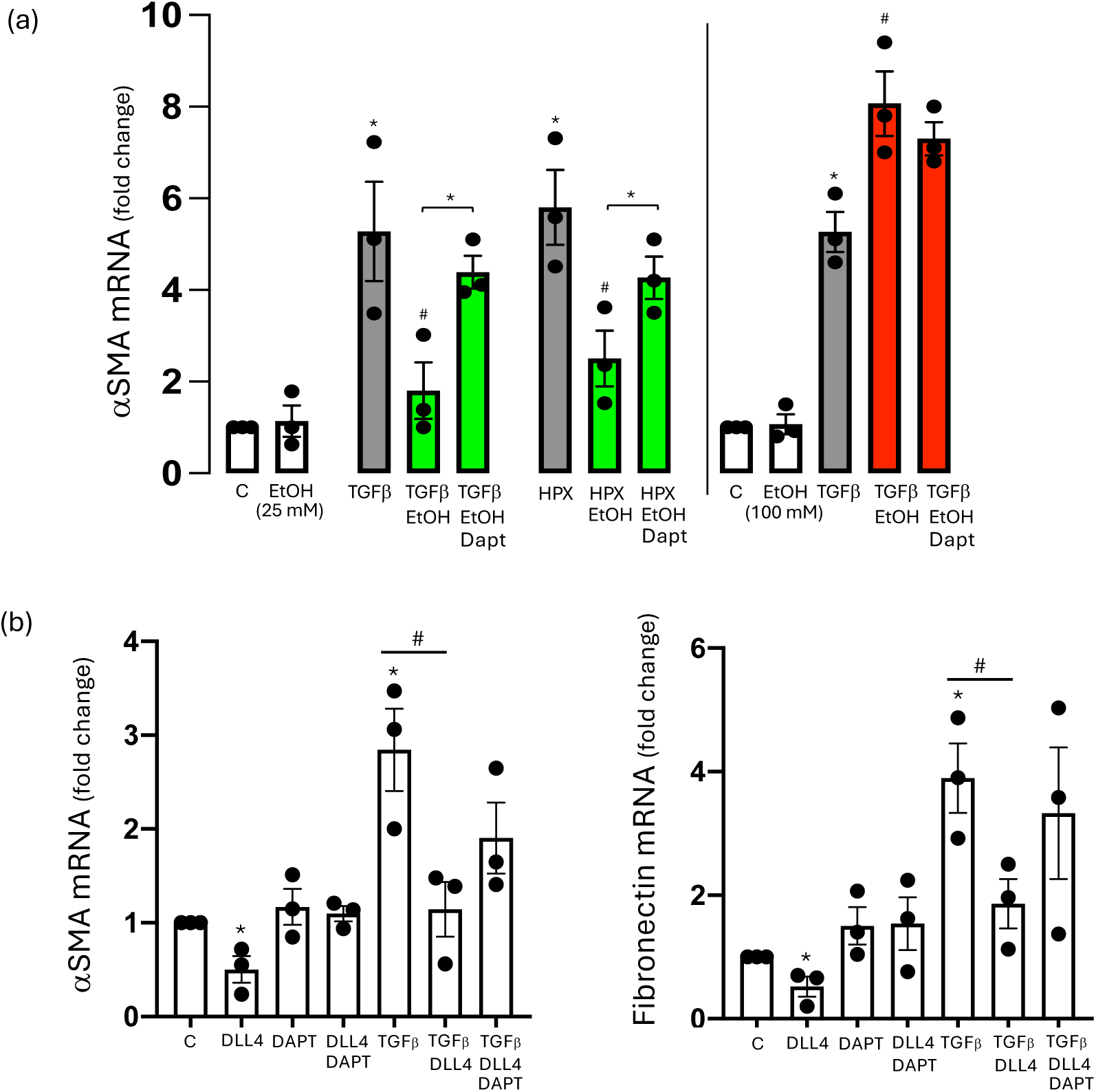
EtOH at moderate, but not high levels, inhibits TGFβ1- and hypoxia-induced EndMT in a Notch-dependent manner. (a) αSMA mRNA levels in HCAEC treated with TGFβ1 (10 ng/ml) or exposed to hypoxic conditions (5% O_2_), in the absence or presence of moderate dose EtOH (25 mM, green bars) or high dose EtOH (100 mM, red bars) +/- the γ-secretase inhibitor DAPT (20 μM). (b) αSMA and fibronectin (FN) mRNA levels in HCAEC treated with TGFβ1 +/- the Notch ligand DLL4 (3 μg/ml) in the absence or presence of DAPT (20 μM). Data are mean±SEM, n=3. *p<0.05 vs control (no treatment), #p<0.05 vs TGFβ1 or hypoxia.

**Fig 4.**
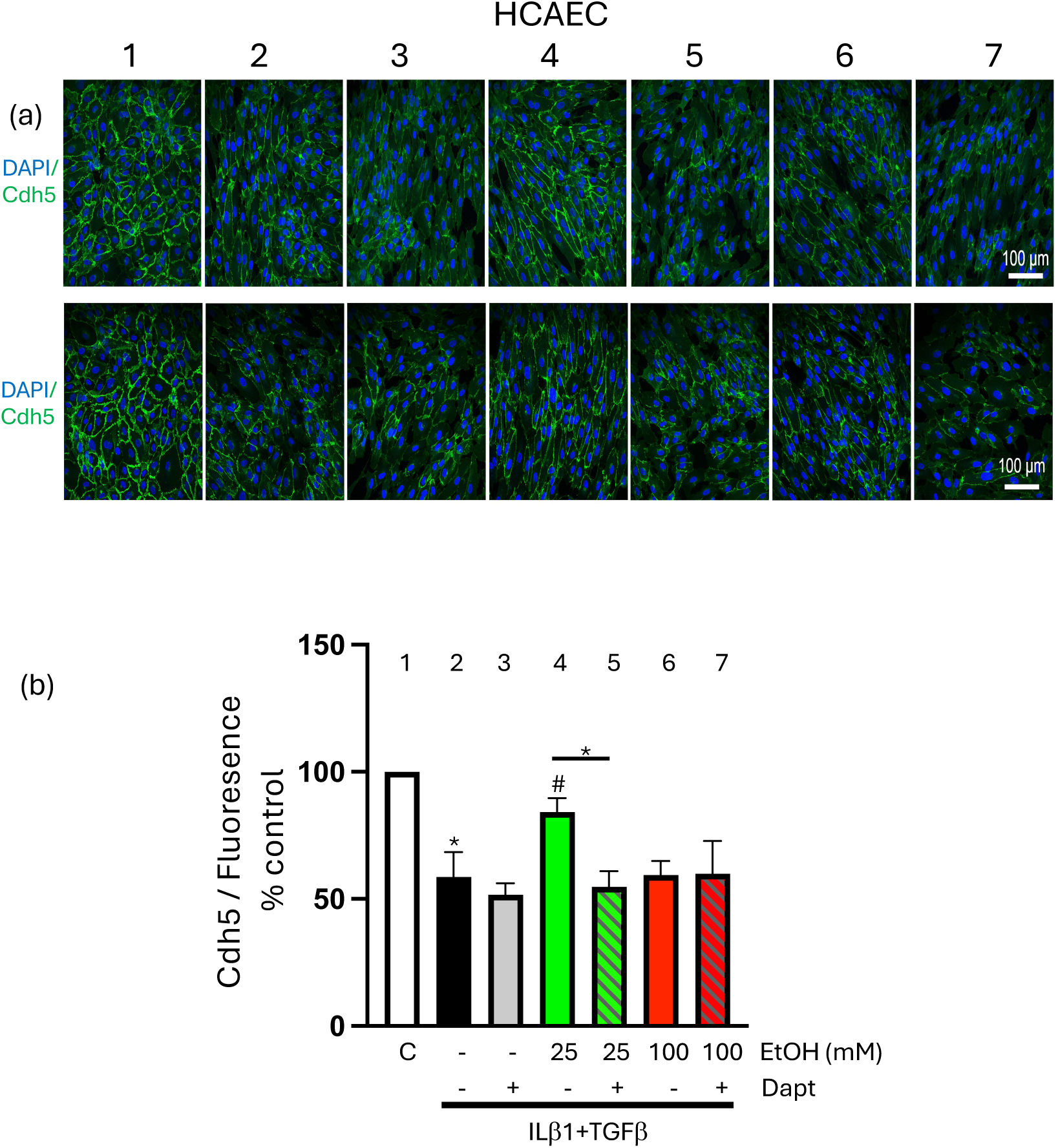
EtOH at moderate, but not high levels, reverses cytokine-induced inhibition of HCAEC Cdh5 expression in a Notch-dependent manner. HCAEC were treated with inflammatory cytokines (IL1β + TGFβ1) in the absence or presence of moderate dose ethanol (25 mM EtOH) or high dose ethanol (100 mM EtOH), with or without the gamma secretase inhibitor DAPT (20 μM), before Cdh5 expression was determined by immunocytochemistry. (a) Representative immunofluorescence images from 2 separate experiments, and (b) cumulative quantitative data of Cdh5 fluorescent staining. Data are mean±SEM, n=6. *p<0.05 vs control (no treatment, grp 1), #p<0.05 vs IL1β + TGFβ1 (grp 2).

### Ethanol dose-dependently inhibits cytokine-stimulated endothelial migration

EndMT is associated with a gain in migratory capacity (1). We assessed migration of endothelial cells by scratch wound assay. Cytokine treatment (IL1β+TGFβ2) stimulated HUVEC migration as indicated by wound closure over time (Fig 5). Co-treatment with moderate dose (25 mM) and with high dose (100 mM) ethanol attenuated the cytokine-induced migration response in a dose-dependent manner (Fig 5).

**Fig 5.**
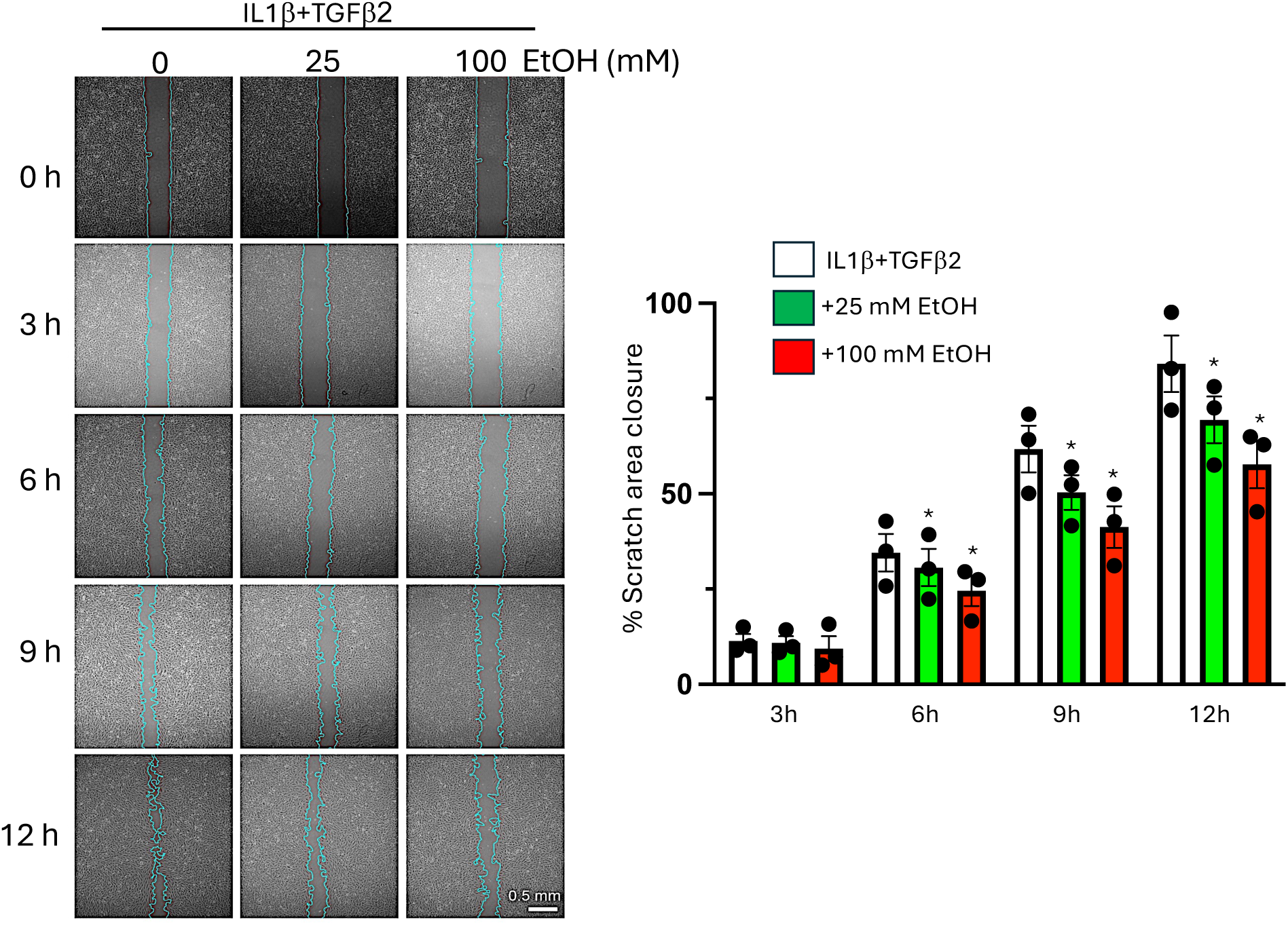
EtOH dose-dependently inhibits cytokine-induced endothelial migration. HUVEC were treated with or without IL1β+TGFβ1 in the absence or presence of moderate (25 mM) and high level (100 mM) EtOH. Migration was determined by scratch wound assay. Representative phase contrast images showing scratch wounds in the different experimental groups at time zero, 3, 6, 9 and 12 h, together with bar graph of cumulative data from 3 separate experiments. Data are mean±SEM, n=3. *p<0.05 vs control (IL1β+TGFβ1 alone).

### Ligation-induced carotid remodeling in Cdh5-Ai9 transgenic mice is attenuated by daily moderate alcohol but aggravated by episodic binge alcohol consumption

To determine the *in vivo* relevance of our cell data, carotid ligation was performed in endothelial lineage-traced Cdh5-Ai9 mice treated with either ‘daily moderate’ EtOH (2 drink equivalent daily) or ‘episodic binge’ EtOH (7 drink equivalent, 2d/wk). No alcohol ‘Controls’ received calorically matched cornstarch solution. Ligation injury induced robust vascular remodeling characterized by adventitial thickening, neointimal hyperplasia, and lumen narrowing (Fig 6, Fig 7), with a more pronounced response in males than in females (Fig 7). Daily moderate ethanol exposure significantly attenuated neointimal formation and preserved lumen area compared with controls, with no significant effect on medial or adventitial compartments (Fig 8). The inhibitory effect of moderate EtOH was greater in males than in females. In contrast, episodic binge ethanol exposure exacerbated adventitial thickening and neointimal hyperplasia relative to controls (Fig 8). These findings demonstrate that alcohol exerts opposing effects on pathologic vascular remodeling depending on the amount and exposure pattern.

**Fig 6.**
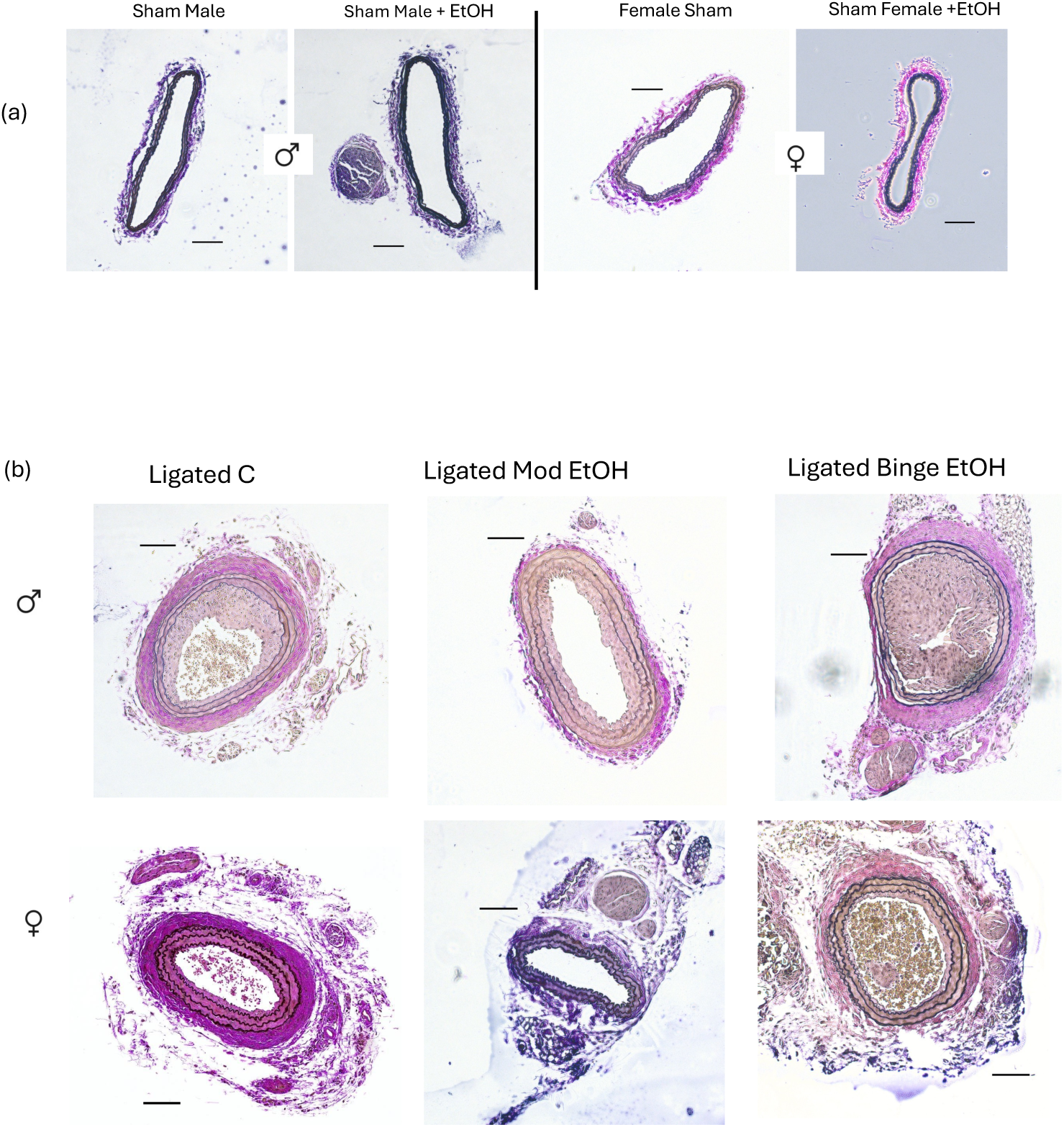
Ligation-induced carotid remodeling. Vessels were harvested 14 days post sham- operation or ligation. (a) Representative Van Gieson stained carotid cross sections from Control and moderate Ethanol (EtOH) sham-operated Cdh5xAi9 male and female mice. (b) Representative Van Gieson stained cross sections from Ligated controls, Ligated + daily moderate EtOH (2 drink equivalent/d, BAC 15 mM), or Ligated + Binge EtOH (7 drink equivalent 2d/wk, BAC 50 mM) cohorts; males upper row, females lower row.

**Fig 7.**
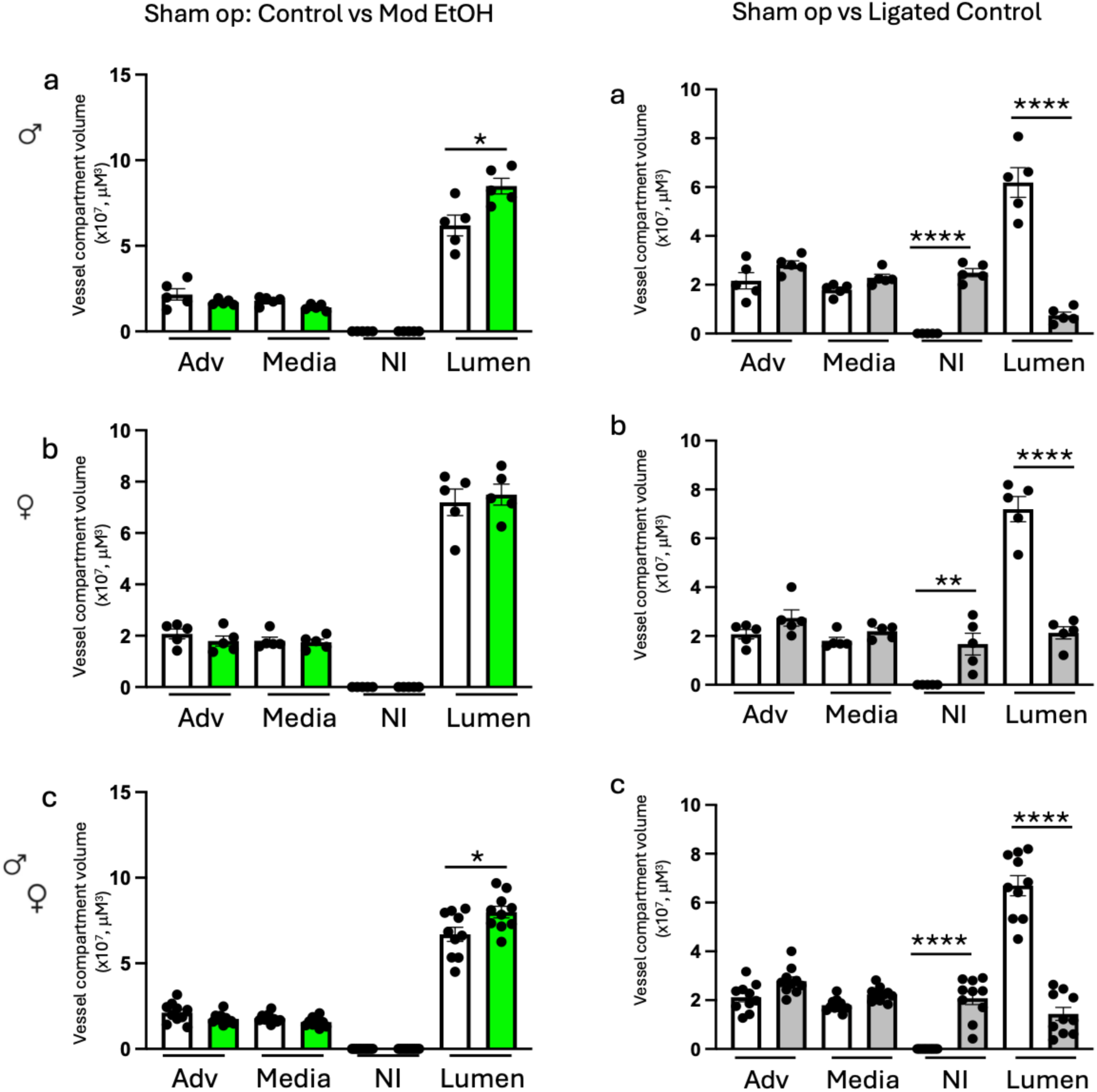
Morphological analysis of carotids from sham and ligated Cdh5xAi9 transgenic mice. (a-c) left hand side; carotid adventitial (adv), medial, neointimal (NI) and lumen volumes in sham- operated controls (white bars) and sham-operated daily moderate EtOH group (green bars). (a- c) right hand side; carotid compartment volumes in sham-operated (white bars) and ligated conytrols (grey bars). Ligation injury stimulated neointimal hyperplasia and decreased lumen volume, to a greater extent in males than females. Data are mean±SEM, n=5 each for males (a) and females (b), (c) n=10, males and females pooled. *p<0.05, ** P<0.001, **** p<0.0001 vs sham-operated control.

**Fig 8.**
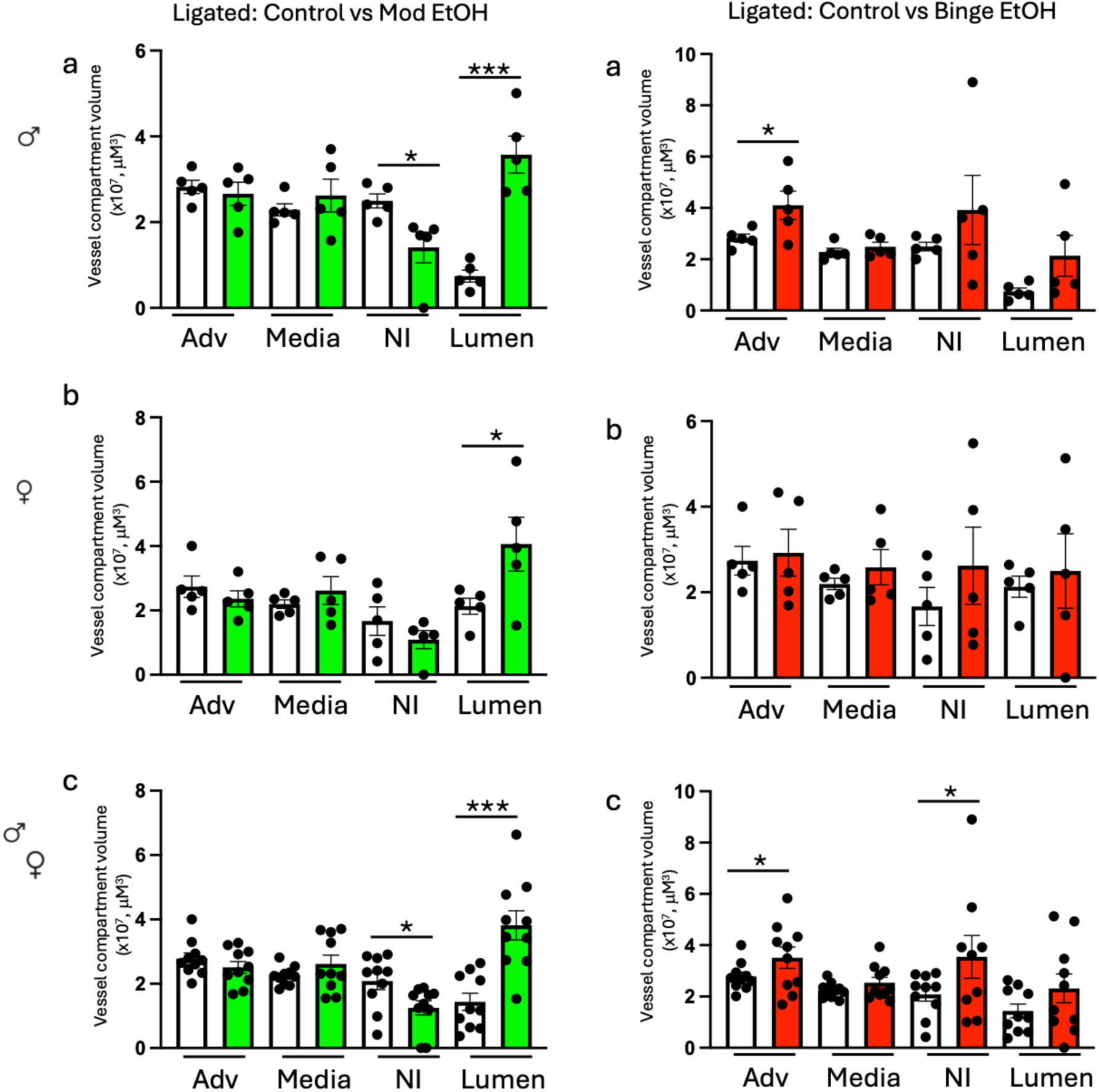
Differential effect of ‘daily moderate’ and ‘episodic binge’ EtOH exposures on ligation-induced carotid remodeling. Daily moderate EtOH exposure (left hand side graphs, green bars) significantly inhibited ligation-induced neointima (NI) formation and increased lumen volume, compared to controls (white bars). In contrast, 2-d binge EtOH exposure (right hand side graphs, red bars) increased adventitial volume and neointima hyperplasia. Data are mean±SEM, n=5 each for males (a) and females (b), (c) n=10, males and females pooled. *p<0.05, ** P<0.001, **** p<0.0001 vs ligated control.

### Differential effect of daily moderate vs episodic binge ethanol on myo-endothelial cell population/EndMT following ligation-injury in mouse carotid

To assess endothelial phenotypic switching *in vivo,* carotid sections were analyzed for cells co-expressing endothelial (Cdh5) and mesenchymal (αSMA) markers i.e., myo-endothelial cells, considered to be indicative of EndMT. In control ligated vessels, a substantial proportion of cells exhibited this dual-marker phenotype: ∼35% of total cells and ∼50% of neointimal cells (Fig 9), consistent with EndMT-like changes. Daily moderate ethanol exposure significantly reduced the proportion and number of these cells, both across the vessel wall and within the neointima (Fig 9). In contrast, episodic binge exposure did not reduce, and in some quantitative analyses significantly increased, the abundance of these cells (Fig 9). These findings are consistent with marked suppression of EndMT-like phenotypic switching by moderate ethanol, but enhancement under binge exposure conditions.

**Fig 9.**
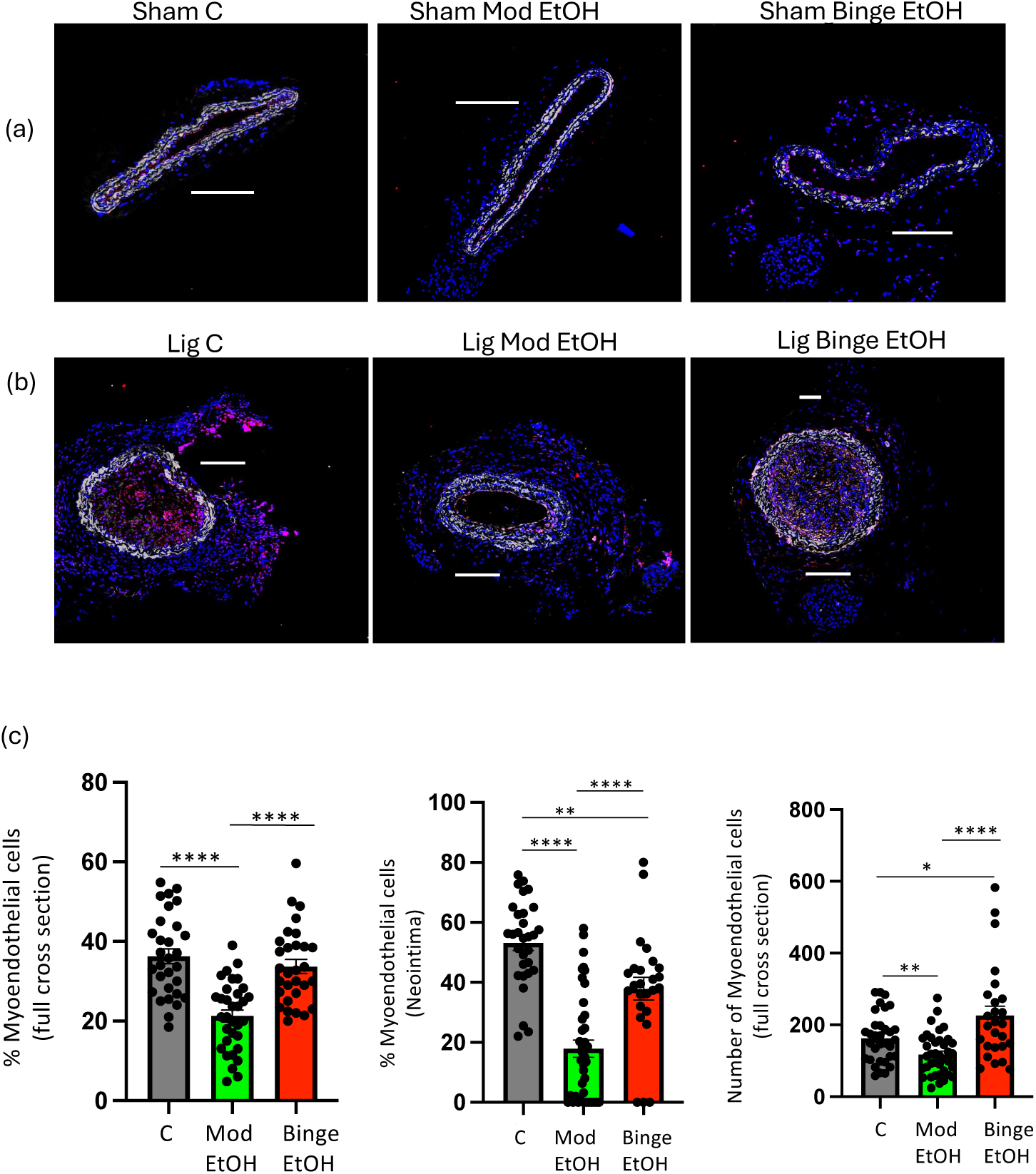
EndMT in remodeled carotids following ligation injury is reduced by moderate EtOH, but not by binge EtOH exposure. (a) (b) Representative images of immunofluorescently stained carotid cross sections from sham-operated and ligated controls, moderate EtOH, and Binge EtOH experimental groups (males). Blue = Dapi nuclear stain, red = Tm-Cdh5+, and white=αSMA+. (c) Cells co-expressing Cdh5 and αSMA (i.e., myo-endothelial cells indicative of EndMT) were quantified using QuPath bioimage analysis software in carotid cross sections post-ligation from controls (grey bars), moderate EtOH (green bars), and binge EtOH (red bars) experimental groups. Bar graphs show cumulative data expressed as % Myoendothelial cells /full cross section, % Myoendothelial cells /neointima, or Number of myoendothelial cells/full cross section. Data are mean±SEM, n=25-30 sections from 5-6 mice). *p<0.05, ** P<0.001, **** p<0.0001.

## DISCUSSION

Here we report for the first time that alcohol exerts a biphasic effect on endothelial plasticity to modulate EndMT and pathologic arterial remodeling. At moderate levels, 1-3 drink equivalent, alcohol inhibited EndMT induced by cytokines and hypoxia *in vitro* and in a carotid ligation injury model *in vivo* concomitant with reduced neointimal hyperplasia, whereas at high levels (7 drink equivalent) alcohol promoted atherogenic endothelial transition and worsened pathologic remodeling. These findings identify regulation of endothelial plasticity as a potential mechanistic link between alcohol consumption patterns and atherosclerotic cardiovascular disease (ASCVD).

Changes in vascular cell fate and plasticity are key drivers of arterial disease (18). Endothelial cells are particularly important in maintaining vessel homeostasis with atherogenesis strongly influenced by altered endothelial behavior (4). One process increasingly recognized as a central contributor to arterial pathology is endothelial-to-mesenchymal transition (EndMT), a phenotypic conversion in which endothelial cells lose their characteristic markers and acquire features typical of smooth muscle–like mesenchymal cells (17, 29). This may be concomitant with a change in morphology, altered extracellular matrix production, and a gain in migratory capacity and contractile function (17). EndMT can be triggered by inflammation, hypoxia, or pathologic flow patterns and contributes to vessel remodeling, neointimal hyperplasia, and vascular fibrosis (5) (32). Thus, the inhibition or reversal of ‘atherogenic EndMT’ can be considered a clinical goal and a therapeutic target of value with respect to atherosclerosis mitigation (32, 34).

The consumption of alcohol is one of several modifiable lifestyle factors that can affect the incidence and severity of ASCVD (8) (24). Ethanol is the type of alcohol found in alcoholic beverages, and the terms ethanol and alcohol are used interchangeably. While the overall health effect of alcohol remains controversial, and while drinking pattern, context (e.g., with /without food), and type of alcohol consumed must be taken into account, there is a considerable body of evidence to support a j-shaped or biphasic effect of alcohol on ASCVD (2). A large number of observational studies and meta analyses, together with some animal and clinical studies, show a lower ASCVD risk for low-to-moderate alcohol consumption (generally considered up to 4 drinks or 60g of alcohol/d in men, and up to ∼3 drinks or 40g alcohol/d in women) compared to abstaining, whereas heavy drinking or bingeing (>4 drinks/d) contributes to increased mortality and greater ASCVD burden (10, 12, 21). Precisely how alcohol at different doses affects arterial cells to ultimately dictate arterial pathology is not fully understood, and no information exists as to whether alcohol affects cell plasticity in this context.

A principal finding of this study is a biphasic effect of alcohol on EndMT. Cytokines and hypoxia induced typical features of EndMT in cultured human endothelial cells including downregulation of classic endothelial markers (eNOS, CD31, Cdh5), and upregulation of mesenchymal markers (αSMA, SM22α), extracellular matrix fibronectin, and the EndMT-associated transcription factor SNAIL, confirming detectable endothelial phenotypic switching under these conditions. Co-treatment of endothelial cells with ‘low-to-moderate’ alcohol doses (5-25 mM) attenuated these changes, whereas higher concentrations (50-100 mM) failed to do so, and in some cases exacerbated the response. Importantly, the inhibitory effect of moderate alcohol was observed across a variety of EndMT-inducing stimuli (inflammatory cytokines alone and in combination, and hypoxia), and in two different cell lines (HCAEC, HUVEC) suggesting a broad modulation by alcohol of endothelial phenotype.

Our data further suggest that Notch signaling contributes, at least in part, to the inhibitory effects of moderate alcohol on EndMT. Pharmacological inhibition of Notch signaling with DAPT abrogated the suppressive effect of moderate alcohol on cytokine- and hypoxia-induced changes in αSMA and Cdh5 expression consistent with involvement of this pathway. These findings are broadly in line with our previous observations demonstrating biphasic regulation of Notch signaling by alcohol in endothelial cells (25, 26), as well as with studies implicating Notch in the maintenance of endothelial homeostasis and suppression of mesenchymal transition (3, 22, 27). However, given the context-dependent roles of Notch signaling in EndMT, including reports supporting both pro- and anti-EndMT effects (19, 31, 33), these findings should be interpreted as supportive rather than definitive. Notably, pro-EndMT effects observed with higher alcohol exposure were not sensitive to Notch inhibition, suggesting that distinct signaling pathways likely mediate the detrimental effects of binge-level exposure.

Functionally, EndMT is associated with a gain in migratory capacity that facilitates cell accumulation and arteriosclerosis (6). Of interest, our data show that ethanol inhibits cytokine-induced migration in a dose-dependent manner, with both moderate and high levels being inhibitory. Another hallmark of EndMT is that the endothelial monolayer becomes leaky having lost cell-cell junctional proteins (1). We have previously reported that alcohol has a biphasic effect on endothelial barrier integrity, with moderate levels improving it but high levels decreasing it (25). Together, these functional findings (migration and barrier integrity) indicate that moderate dose alcohol preserves endothelial phenotype to a greater extent than high level alcohol exposure, limiting acquisition of the migratory, mesenchymal phenotype and preserving barrier function. In the context of vascular injury, such effects would be expected to limit endothelial contribution to neointimal cell populations, thereby reducing pathological remodeling.

The *in vivo* data provide important physiological context for these observations and further support the inhibitory effects of moderate, but not high dose alcohol exposure, on EndMT. In the carotid ligation model, daily moderate alcohol significantly reduced neointimal hyperplasia and preserved lumen diameter, whereas episodic binge exposure exacerbated pathological remodeling. These opposite structural changes were paralleled by alterations in the abundance of myo-endothelial cells. Moderate alcohol reduced, whereas binge exposure increased, this cell population within the remodeled vessel wall. While co-expression of endothelial and mesenchymal markers is widely used as an indicator of EndMT, we acknowledge that definitive lineage conversion cannot be established without higher-resolution approaches such as single-cell trajectory analysis or lineage tracing at the transcriptomic level. Nonetheless, the concordance between *in vitro* phenotypic modulation and *in vivo* remodeling and myoendothelial population emergence strongly supports a role for endothelial plasticity in mediating these effects. These findings provide a mechanistic framework linking alcohol exposure patterns to vascular disease risk. Our data suggest that differential regulation of endothelial plasticity, specifically EndMT, may explain at least in part, the frequently reported j-shaped relationship between alcohol consumption and ASCVD.

In summary, our data demonstrate that alcohol modulates endothelial phenotypic plasticity in a biphasic manner. Low-to-moderate dose alcohol suppresses EndMT to maintain endothelial phenotype and attenuate pathologic arterial remodeling, whereas high dose alcohol fails to confer these atheroprotective effects and instead promotes EndMT and neointimal hyperplasia. These findings identify EndMT as a key cellular process targeted by alcohol in a hormetic manner to modulate vascular pathology and ASCVD, providing new mechanistic insight into the divergent health consequences associated with different exposure levels and patterns of alcohol drinking.

## Data availability

All data included in this study are available upon reasonable request from the corresponding author (EMR). This study does not generate any unique reagents or original code. Requests for further information should be directed to Eileen M Redmond.

## Acknowledgements.

We thank Diana Scott for histological tissue processing, Conor Vanderlip for carotid image cataloging, and Dr Kaye Thomas for confocal microscopic expertise.

## Grants

This work was supported by grants RO1AA024082 and RO1AA031225 (to EMR) from the National Institutes of Health.

## Disclosures

No conflicts of interest, financial or otherwise, are declared by the authors.

## Author contributions

PAC and EMR conceived and designed research; WL, YG, FA and NR performed experiments; WL, YG, FA, NR and EMR analyzed data; WL, YG, FA, NR, PAC and EMR interpreted results of experiments; YG, FA, NR and EMR prepared figures; EMR drafted manuscript; PAC and EMR edited and revised manuscript; WL, YG, FA, NR, PAC and EMR approved final version of manuscript.

## Notes

### Competing Interest Statement

The authors have declared no competing interest.

## References

1. Alvandi Z and Bischoff J. Endothelial-Mesenchymal Transition in Cardiovascular Disease. Arteriosclerosis, thrombosis, and vascular biology 41: 2357–2369, 2021.

2. Alvarez-Mon MA, Martinez-Urbistondo D, Barberia-Latasa M, Vazquez-Ruiz Z, Ruiz-Canela M, Bes-Rastrollo M, and Martinez-Gonzalez MA. The Unfinished Debate on Wine and Other Alcoholic Beverages: Conflicting Evidence, Public Health Messages and the Missing Trial. Nutrients 18, 2026.

3. Aquila G, Kostina A, Vieceli Dalla Sega F, Shlyakhto E, Kostareva A, Marracino L, Ferrari R, Rizzo P, and Malaschicheva A. The Notch pathway: a novel therapeutic target for cardiovascular diseases? Expert Opin Ther Targets 23: 695–710, 2019.

4. Cahill PA and Redmond EM. Vascular endothelium - Gatekeeper of vessel health. Atherosclerosis 248: 97–109, 2016.

5. Chang CJ, Lai YJ, Tung YC, Wu LS, Hsu LA, Tseng CN, Chang GJ, Yang KC, and Yeh YH. Osteopontin mediation of disturbed flow-induced endothelial mesenchymal transition through CD44 is a novel mechanism of neointimal hyperplasia in arteriovenous fistulae for hemodialysis access. Kidney Int 103: 702–718, 2023.

6. Chen PY, Schwartz MA, and Simons M. Endothelial-to-Mesenchymal Transition, Vascular Inflammation, and Atherosclerosis. Front Cardiovasc Med 7: 53, 2020.

7. Colpani V, Baena CP, Jaspers L, van Dijk GM, Farajzadegan Z, Dhana K, Tielemans MJ, Voortman T, Freak-Poli R, Veloso GGV, Chowdhury R, Kavousi M, Muka T, and Franco OH. Lifestyle factors, cardiovascular disease and all-cause mortality in middle-aged and elderly women: a systematic review and meta-analysis. Eur J Epidemiol 33: 831–845, 2018.

8. Costanzo S, Di Castelnuovo A, Donati MB, Iacoviello L, and de Gaetano G. Alcohol consumption and mortality in patients with cardiovascular disease: a meta-analysis. Journal of the American College of Cardiology 55: 1339–1347, 2010.

9. de Queiroz DB, Parente JM, Pernomian L, Waigi EW, Alfaidi M, Tan W, McCarthy CG, and Wenceslau CF. Endothelial Cell Phenotypic Plasticity in Cardiovascular Physiology and Disease: Mechanisms and Therapeutic Prospects. Am J Hypertens 38: 411–421, 2025.

10. Di Castelnuovo A, Costanzo S, Bagnardi V, Donati MB, Iacoviello L, and de Gaetano G. Alcohol dosing and total mortality in men and women: an updated meta-analysis of 34 prospective studies. Archives of internal medicine 166: 2437–2445, 2006.

11. Evrard SM, Lecce L, Michelis KC, Nomura-Kitabayashi A, Pandey G, Purushothaman KR, d’Escamard V, Li JR, Hadri L, Fujitani K, Moreno PR, Benard L, Rimmele P, Cohain A, Mecham B, Randolph GJ, Nabel EG, Hajjar R, Fuster V, Boehm M, and Kovacic JC. Endothelial to mesenchymal transition is common in atherosclerotic lesions and is associated with plaque instability. Nat Commun 7: 11853, 2016.

12. Fernandez-Sola J. Cardiovascular risks and benefits of moderate and heavy alcohol consumption. Nat Rev Cardiol 12: 576–587, 2015.

13. Fitzpatrick E, Han X, Liu W, Corcoran E, Burtenshaw D, Morrow D, Helt JC, Cahill PA, and Redmond EM. Alcohol Reduces Arterial Remodeling by Inhibiting Sonic Hedgehog-Stimulated Stem Cell Antigen-1 Positive Progenitor Stem Cell Expansion. Alcoholism, clinical and experimental research 41: 2051–2065, 2017.

14. Gusti Y, Liu W, Athar F, Cahill PA, and Redmond EM. Endothelial Homeostasis Under the Influence of Alcohol-Relevance to Atherosclerotic Cardiovascular Disease. Nutrients 17, 2025.

15. Hendriks HFJ. Alcohol and Human Health: What Is the Evidence? Annu Rev Food Sci Technol 11: 1–21, 2020.

16. Hong L, Du X, Li W, Mao Y, Sun L, and Li X. EndMT: A promising and controversial field. Eur J Cell Biol 97: 493–500, 2018.

17. Kovacic JC, Dimmeler S, Harvey RP, Finkel T, Aikawa E, Krenning G, and Baker AH. Endothelial to Mesenchymal Transition in Cardiovascular Disease: JACC State-of-the-Art Review. J Am Coll Cardiol 73: 190–209, 2019.

18. Lin A, Miano JM, Fisher EA, and Misra A. Chronic inflammation and vascular cell plasticity in atherosclerosis. Nat Cardiovasc Res 3: 1408–1423, 2024.

19. Lin QQ, Zhao J, Zheng CG, and Chun J. Roles of notch signaling pathway and endothelial-mesenchymal transition in vascular endothelial dysfunction and atherosclerosis. Eur Rev Med Pharmacol Sci 22: 6485–6491, 2018.

20. Liu W, Harman S, DiLuca M, Burtenshaw D, Corcoran E, Cahill PA, and Redmond EM. Moderate Alcohol Consumption Targets S100beta(+) Vascular Stem Cells and Attenuates Injury-Induced Neointimal Hyperplasia. Alcoholism, clinical and experimental research 44: 1734–1746, 2020.

21. Liu W, Redmond EM, Morrow D, and Cullen JP. Differential effects of daily-moderate versus weekend-binge alcohol consumption on atherosclerotic plaque development in mice. Atherosclerosis 219: 448–454, 2011.

22. Mack JJ and Iruela-Arispe ML. NOTCH regulation of the endothelial cell phenotype. Current opinion in hematology 25: 212–218, 2018.

23. Morrow D, Cullen JP, Liu W, Cahill PA, and Redmond EM. Alcohol inhibits smooth muscle cell proliferation via regulation of the Notch signaling pathway. Arteriosclerosis, thrombosis, and vascular biology 30: 2597–2603, 2010.

24. Piano MR, Marcus GM, Aycock DM, Buckman J, Hwang CL, Larsson SC, Mukamal KJ, Roerecke M, on behalf the American Heart Association Council on L, Cardiometabolic H, Council on C, Stroke N, Council on Clinical C, and Stroke C. Alcohol Use and Cardiovascular Disease: A Scientific Statement From the American Heart Association. Circulation 152: e7–e21, 2025.

25. Rajendran NK, Liu W, Cahill PA, and Redmond EM. Alcohol and vascular endothelial function: Biphasic effect highlights the importance of dose. Alcohol Clin Exp Res (Hoboken) 47: 1467–1477, 2023.

26. Rajendran NK, Liu W, Cahill PA, and Redmond EM. Caveolin-1 inhibition mediates the opposing effects of alcohol on gamma-secretase activity in arterial endothelial and smooth muscle cells. Physiol Rep 11: e15544, 2023.

27. Rajendran NK, Liu W, Chu CC, Cahill PA, and Redmond EM. Moderate dose alcohol protects against serum amyloid protein A1-induced endothelial dysfunction via both notch-dependent and notch-independent pathways. Alcoholism, clinical and experimental research 45: 2217–2230, 2021.

28. Ronksley PE, Brien SE, Turner BJ, Mukamal KJ, and Ghali WA. Association of alcohol consumption with selected cardiovascular disease outcomes: a systematic review and meta-analysis. BMJ (Clinical research ed 342: d671, 2011.

29. Souilhol C, Harmsen MC, Evans PC, and Krenning G. Endothelial-mesenchymal transition in atherosclerosis. Cardiovascular research 114: 565–577, 2018.

30. Suarez-Arnedo A, Torres Figueroa F, Clavijo C, Arbelaez P, Cruz JC, and Munoz- Camargo C. An image J plugin for the high throughput image analysis of in vitro scratch wound healing assays. PLoS One 15: e0232565, 2020.

31. Tian D, Zeng X, Wang W, Wang Z, Zhang Y, and Wang Y. Protective effect of rapamycin on endothelial-to-mesenchymal transition in HUVECs through the Notch signaling pathway. Vascul Pharmacol 113: 20–26, 2019.

32. Xu Y, Korayem A, Cruz-Solbes AS, Chandel N, Sakata T, Mazurek R, Mavropoulos SA, Kariya T, Aikawa T, Yamada KP, D’Escamard V, V’Gangula B, Baker AH, Ma L, Bjorkegren JLM, Fuster V, Boehm M, Fish KM, Tadros R, Ishikawa K, and Kovacic JC. Inhibition of endothelial-to-mesenchymal transition in a large animal preclinical arteriovenous fistula model leads to improved remodelling and reduced stenosis. Cardiovascular research 120: 1768–1779, 2024.

33. Yang R, Yang F, Hu Y, Chen M, Liu Y, Li J, and Zhong W. Hepatocyte Growth Factor Attenuates the Development of TGF-beta1- Induced EndMT through Down-regulating the Notch Signaling. Endocr Metab Immune Disord Drug Targets 20: 781–787, 2020.

34. You N, Liu G, Yu M, Chen W, Fei X, Sun T, Han M, Qin Z, Wei Z, and Wang D. Reconceptualizing Endothelial-to-mesenchymal transition in atherosclerosis: Signaling pathways and prospective targeting strategies. J Adv Res 77: 419–441, 2025.

35. Zhang H, Lui KO, and Zhou B. Endocardial Cell Plasticity in Cardiac Development, Diseases and Regeneration. Circulation research 122: 774–789, 2018

